# Parallel accumulation – serial fragmentation combined with data-independent acquisition (diaPASEF): Bottom-up proteomics with near optimal ion usage

**DOI:** 10.1101/656207

**Authors:** Florian Meier, Andreas-David Brunner, Max Frank, Annie Ha, Isabell Bludau, Eugenia Voytik, Stephanie Kaspar-Schoenefeld, Markus Lubeck, Oliver Raether, Ruedi Aebersold, Ben C. Collins, Hannes L. Röst, Matthias Mann

**Affiliations:** Proteomics and Signal Transduction, Max Planck Institute of Biochemistry, Martinsried, Germany; Donnelly Centre for Cellular and Biomolecular Research, University of Toronto, Toronto, Canada; Bruker Daltonik GmbH, Bremen, Germany; Department of Biology, Institute of Molecular Systems Biology, ETH Zurich, Zurich, Switzerland; Faculty of Science, University of Zurich, Zurich, Switzerland; School of Biological Sciences, Queen’s University of Belfast, UK; NNF Center for Protein Research, University of Copenhagen, Copenhagen, Denmark

## Abstract

Data independent acquisition (DIA) modes isolate and concurrently fragment populations of different precursors by cycling through segments of a predefined precursor *m/z* range. Although these selection windows collectively cover the entire *m/z* range, overall only a few percent of all incoming ions are sampled. Making use of the correlation of molecular weight and ion mobility in a trapped ion mobility device (timsTOF Pro), we here devise a novel scan mode that samples up to 100% of the peptide precursor ion current. We extend an established targeted data extraction workflow by including the ion mobility dimension for both signal extraction and scoring, thereby increasing the specificity for precursor identification. Data acquired from whole proteome digests and mixed organism samples demonstrate deep proteome coverage and a very high degree of reproducibility as well as quantitative accuracy, even from 10 ng sample amounts.

## INTRODUCTION

Mass spectrometry (MS)-based proteomics, like other omics technologies, aims for an unbiased, comprehensive and quantitative counteracting electric field. Trapped ions description of the system under investigation^1–3^. Proteomics workflows have become increasingly successful in characterizing complex proteomes in great depth^4,5^. For the application of this technology to large sample cohorts e.g. for systematic screening or clinical applications, which require a high degree of reproducibility and data completeness, data independent acquisition (DIA) schemes are particularly attractive^6,7^. Unlike in data dependent acquisition (DDA) where particular precursors are sequentially fragmentation. PASEF achieves a more selected, in DIA groups of ions are recursively isolated by the quadrupole and concurrently fragmented, thus generating convoluted fragment ion spectra composed of fragments from many different precursors^8–10^. Although DIA guarantees that each precursor in a predefined mass range is fragmented once per cycle, spectral complexity poses a great challenge to subsequent analysis^11^. This is reduced by narrow isolation windows, but these increase the cycle times needed to cover the entire mass range. Moreover, as every precursor is only isolated once per cycle, the ion sampling efficiency at the mass selective quadrupole for DIA methods is limited to 1-3% with typical schemes of 32 or 64 windows.

Adding ion mobility separation to the chromatographic and mass separation should increase sensitivity and reduce spectral complexity^12–15^. The trapped ion mobility spectrometer (TIMS) is a particularly compact mobility analyzer in which ions are captured in an RF ion tunnel by the opposing forces of the gas flow from the source and the ^16–18^. Trapped ions are then sequentially released as a function of their collisional cross section by lowering the electric potential. Mobility resolution depends on the ramp time, which is typically 50 to 100 ms, a time range between chromatographic peak widths (seconds) and the time-of-flight (TOF) spectral acquisition (about 100 μs per pulse). In a TIMS-quadrupole-TOF configuration, the release of precursor ions can be synchronized with the quadrupole selection in a method termed parallel accumulation followed by serial fragmentation^19^. PASEF achieves a more than ten-fold increase in sequencing speed in data dependent acquisition, without the loss of sensitivity that is otherwise inherent to very fast fragmentation cycles^20,21^.

Here we investigate if the PASEF principle can be extended to DIA, combining the advantages of this acquisition method with the inherent efficiency of PASEF. To realize this vision, we modified the mass spectrometer to support ‘diaPASEF’ acquisition cycles. Building on open-source software^22^ we perform targeted extraction of fragment ion traces from the four dimensional data space to confirm the identity and indicate the abundance of the peptides in the sample. We explore the performance of the diaPASEF principle in typical proteomics applications such as single run proteome analysis, label-free quantification, as well as the in-depth characterization of extremely low sample amounts.

## RESULTS

### The diaPASEF principle

In the timsTOF Pro instrument (Bruker Daltonik), peptides separated by liquid chromatography are ionized, introduced into the mass spectrometer and immediately trapped in a first TIMS device (TIMS1, **Fig. 1a**). They are then transferred into TIMS2 from which they are released in reversed order of their ion mobilities (largest ions released first). In parallel, incoming ions are again accumulated in TIMS1, assuring full ion utilization. If operated in MS1 mode, the ion species sequentially released from TIMS2 reach the orthogonal accelerator from which rapid TOF pulses result in high-resolution mass spectra (>35,000 over the entire mass range). If operated in MS/MS mode, specific precursors or groups of precursors are selected by a quadrupole and transferred to a collision cell. The *m/z* of the resulting fragment ions are then analyzed at full resolution and speed (about 10 kHz) in the TOF analyzer. For peptide ions of a given charge state, ion mobilities and precursor masses are correlated (**Fig. 1b**). We reasoned that this feature could be used to isolate precursor mass windows for DIA without losing the ions outside of the respective windows as is the case in other DIA acquisition schemes. Because high *m/z* (low ion mobility) ions are stored at the end of the voltage gradient, they are released first and the mass selective quadrupole therefore needs to be first positioned at high *m/z*. Coordinately with the decreasing *m/z* of the ions sequentially released from the TIMS, the quadrupole mass isolation window slides down to lower *m/z* values such that the trapped ion cloud is fully transmitted (grey area in **Fig. 1c**). To approximate this ideal diaPASEF scan, we stepped the isolation window as a function of TIMS release time, covering the vast majority of precursors of the 2^+^ and 3^+^ charge state (**Fig. 1d**). Implementation of this principle required novel firmware able to synchronize collision energies with the mass selection (**Methods**). Subsequent fragmentation in the collision cell distributes the fragments of each DIA window at the exact ion mobility position of the precursor (**Fig. 1e**). Over the chromatographic elution of a precursor, the intensities of its fragments follow the precursor intensity in time (z-direction). The signal traced out by the set of fragments of an individual precursor is a set of very flat ellipsoids (x or *m/z* dimension), spreading in ion mobility direction (y-direction) and elongated in the retention time dimension (z-dimension). For the entire liquid chromatography tandem mass spectrometry (LC-MS/MS) run, this leads to a ‘perfect data cuboid’ in four dimensional space, containing all fragment ions of all precursors over the entire elution time, with signal intensity as the fourth dimension.

**Figure 1.**
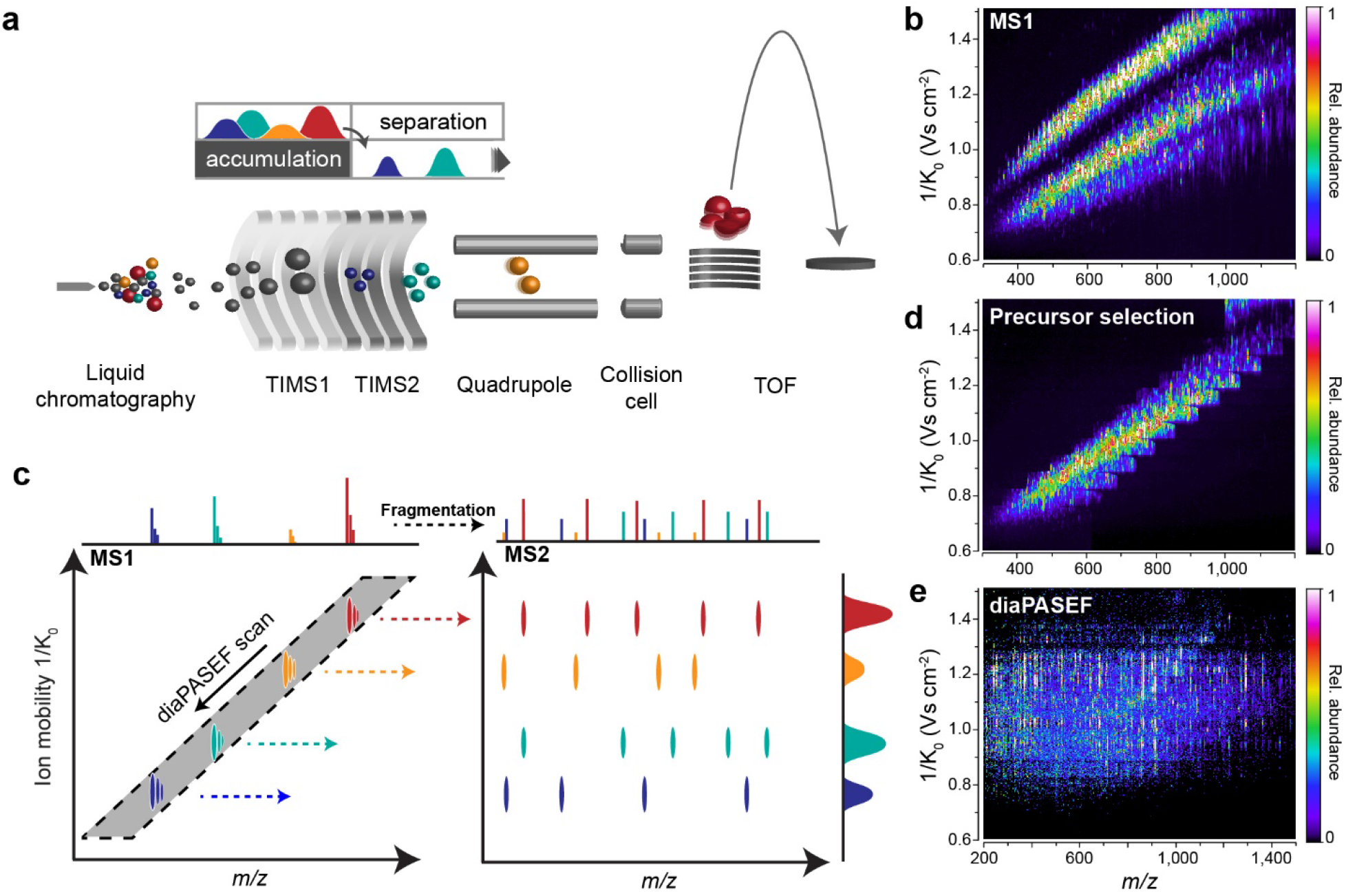
The diaPASEF acquisition method. **a**, Schematic ion path of the timsTOF Pro mass spectrometer. **b**, Correlation of peptide ion mobility and *m/z* in a tryptic digest of HeLa cell lysate. **c**, In diaPASEF, the quadrupole isolation window (grey) is dynamically positioned as a function of ion mobility (arrow). In a single TIMS scan, ions from the selected mass ranges are fragmented to record ion mobility-resolved MS2 spectra of all precursors. **d**, Implementation of diaPASEF precursor selection with a stepped quadrupole isolation scheme. **e**, Representative example of a single diaPASEF scan with the precursor selection scheme from d.

### Quantifying the increase in ion sampling efficiency

To explore the diaPASEF principle in practice, we measured a tryptic digest of bovine serum albumin (BSA) and compared the signals obtained across DDA, DIA and diaPASEF acquisition methods. As a typical example, the peptide DLGEEHFK eluted over 9 s (**Fig. 2a**). In DDA, the doubly charged precursor was accumulated before fragmentation once for 100 ms at the beginning of the elution peak. This corresponds to about 1% of the total elution time and much less than 1% of the entire precursor ion population, as estimated by the relative peak area. In DIA, with a comparably fast cycle time of 1.6 s, the peptide was fragmented seven times over its elution profile, which is sufficient to reconstruct the chromatographic peak shape. In that scheme only a small proportion of the total ion signal (less than 5%) was captured. In contrast, the diaPASEF scheme (**Suppl. Fig. 1**) sampled the fragments in each TIMS scan and for a total of over 100 times, resulting in a nearly complete record of the fragments at every time point. Total efficiency in terms of acquisition time was 96% (because of the interspersed full scans), approaching 100%.

**Figure 2.**
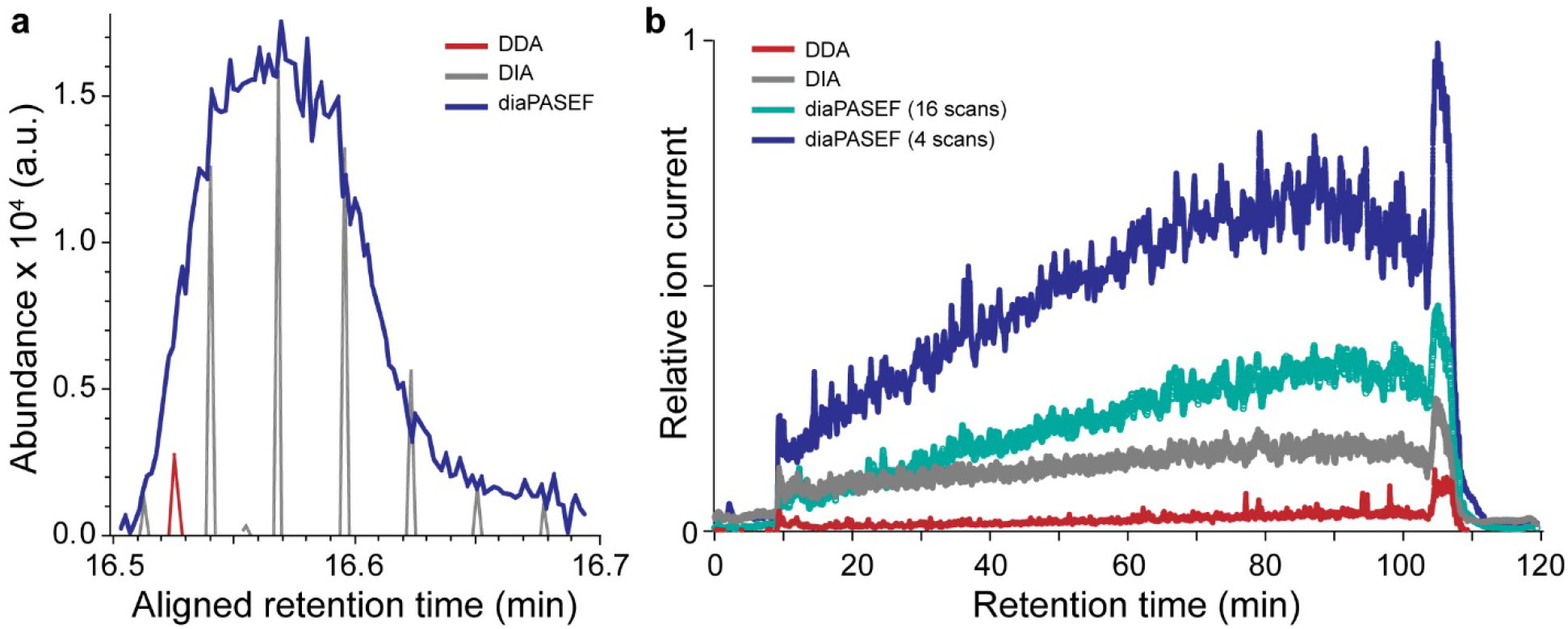
Efficiency of different data acquisition methods. **a**, Extracted fragment ion chromatogram of the y6 ion of the doubly charged DLGEEHFK peptide precursor in an LC-MS analysis of bovine serum album digest acquired with typical DDA and DIA methods as well as a 100% duty cycle diaPASEF method. **b**, Detected ion current from multiply charged precursors in single run analyses of a HeLa digest acquired with DDA, DIA and two diaPASEF schemes. To extract the ion current after quadrupole isolation, no collision energy was applied and the ion current was summed for each TIMS scan. The plot shows the rolling average of 60 TIMS scans.

We next studied the ion sampling efficiency for a HeLa cell tryptic digest. To address the very high density of fragment ions in the data cuboid, our diaPASEF scheme can balance the width of the dynamic isolation window in the m/z dimension (reducing fragment spectral complexity) with the number of TIMS ramps needed to cover the entire precursor space (reducing duty cycle). To exemplify, we here chose a scheme with only four diaPASEF scans, each isolating about 1/4^th^ of all precursors with 50 Th isolation windows, and another scheme with 16 diaPASEF scans and 25 Th isolation windows (**Supplementary Figs. 2 and 3**).

To assess the relative performance of these schemes, we performed DDA and DIA measurements on aliquots of the same sample, using typical parameters for either method. We applied no collision energy and extracted the total ion current of all isolated precursors in the expected peptide space in the *m/z*-ion mobility plane. For the DIA measurements the sampled fraction of the ion current was about three-fold higher than for DDA scans, whereas the four-ramp diaPASEF scheme further increased the accumulated peptide ion current by a factor of five compared to the DIA scan (**Fig. 2b**). We conclude that the diaPASEF principle yields the expected increase in data acquisition efficiency in both simple and complex proteomes.

### Targeted data extraction in four dimensions

To identify and quantify peptides from this novel data structure, we developed Mobi-DIK (Ion Mobility DIA Analysis Kit), a software capable of analyzing the four-dimensional diaPASEF data space (**Fig. 3a**). The workflow is based on the targeted extraction of sets of fragment ions of a specific precursor from the acquired dataset over chromatographic elution time, followed by statistical scoring of the generated peak group with regard to a spectral library. Mobi-DIK extends the targeted data analysis principle for DIA data^23^, as implemented in the OpenSWATH software suite^22^ to the fourth data dimension generated by diaPASEF. The workflow first generates ion mobility-enabled spectral libraries directly from data-dependent PASEF runs using, for instance, the MaxQuant^24,25^ output and stores them in the PQP format. The spectral library is processed using OpenMS tools^26,27^, which we extended to support ion mobility. Calibration between the assay library and experimental DIA data is automatically performed in the *m/z*, retention time and ion mobility dimensions, based on a user-defined set of high confidence peptides (**Methods**). The Mobi-DIK package employs the Bruker interface to query native .tdf files and converts them to mzML files using the vendor API to assign the multiple quadrupole isolation positions in diaPASEF data to individual TOF scans. The algorithm then uses the targeted extraction paradigm

**Figure 3.**
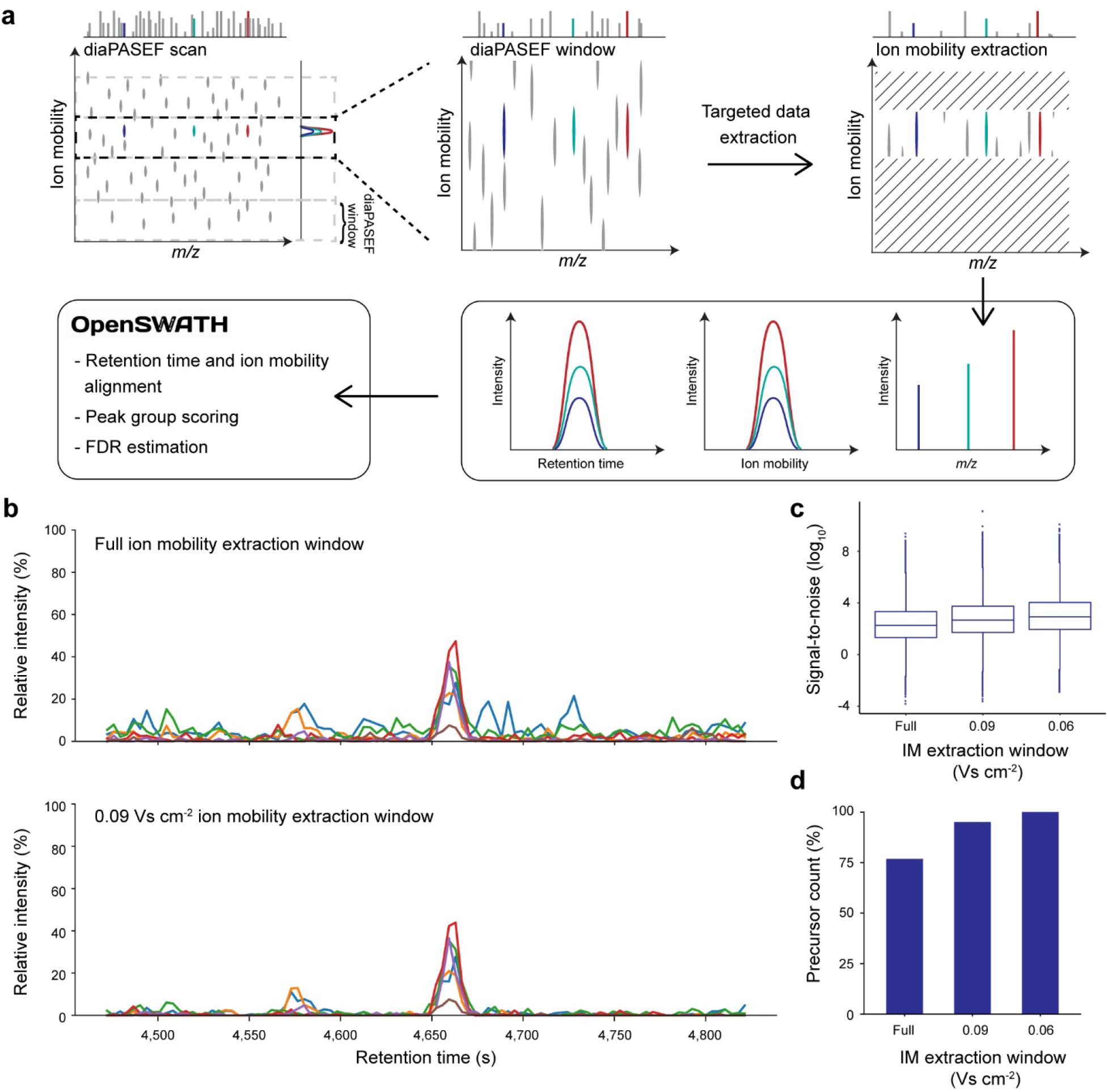
Ion mobility-aware targeted data extraction. **a**, Steps in the Mobi-DIK workflow to extract fragment ion chromatograms from diaPASEF scans. First, our workflow de-multiplexes diaPASEF data into individual diaPASEF windows and then uses the assay library coordinates to perform targeted extraction in the 4-dimensional analysis space of ion mobility, retention time and *m/z*. For each peptide precursor, a full extracted ion chromatogram, as well as an accompanying ion mobilogram and a set of high resolution MS2 spectra are extracted from the data and used for scoring. The OpenSWATH algorithm is used to perform peak group scoring, FDR estimation and alignment using the ion mobility enhanced data. **b**, Example chromatograms extracted at with and without restriction in the ion mobility dimension. **c,** Narrow extraction in ion mobility space improves signal-to-noise (S/N) and removes interfering signals from co-eluting precursors in the same diaPASEF window. The mean S/N ratio increases 2.6-to 4-fold when comparing full ion mobility extraction to narrow extraction. Boxplots show the median (center line), 25^th^ and 75^th^ percentile (lower and upper box limits), the 1.5 × the interquartile range (whiskers) and outliers (points). **d,** Relative number of detected peptide precursors at 1% FDR as a function of the ion mobility extraction window.

for DIA data to construct four-dimensional data cuboids with a user-defined width in *m/z* (ppm), retention time (s) and ion mobility (Vs cm^−2^). These are projected onto the retention time and ion mobility axes to obtain fragment ion chromatograms and mobilograms for each precursor-to-fragment transition in the spectral library. Restricting the extraction data cube in ion mobility dimension to a user-defined width improves the signal-to-noise by removing signals from co-eluting peptide species in the very same precursor mass window, as exemplified in **Fig. 3b**. Investigating the signal-to-noise ratios of all transitions in a single run analysis of HeLa digest, we found that narrowing the ion mobility extraction window to 0.06 Vs cm^−2^, resulting in an average 4-fold increase in signal-to-noise (**Fig. 3c**). Note that the positioning of the quadrupole in diaPASEF already removes interfering ions with very different ion mobility such as singly charged species, therefore the true gain in signal-to-noise as compared with the respective value of a DIA experiment without ion mobility is even higher.

From the projected precursor-to-fragment transitions traces, we next pick peak groups along the chromatographic dimension using the OpenSWATH peak picking and scoring modules. This step in the data analysis workflow selects putative peak candidates that are subsequently scored based on their chromatographic co-elution, goodness of library match and correlation with the precursor profile^22^. For Mobi-DIK, we extended these modules to include scoring along the ion mobility dimension. We make use of the high precision of TIMS ion mobility measurements (<1% in replicates of complex samples^20^) to compute a discriminatory score based on the difference between the library-recorded and the calibrated experimental ion mobility that is combined with the other scores described above. Furthermore, we additionally extract full ion mobilograms for each fragment ion to score the mobility peak shape as well as the peak consistency between all fragment ions. In the single-run analysis of a whole-cell HeLa digest (see below), targeted extraction in the ion mobility dimension (combined with ion mobility-aware scoring) increased peptide identifications compared to a naive analysis by more than 25% (**Fig. 3d**).

### Single run proteome analysis

Having established the diaPASEF acquisition method and the Mobi-DIK data analysis workflow, we next investigated single-run proteome analysis of a human HeLa cancer cell line. First, we built a project-specific library from 24 high-pH reversed-phase peptide fractions with data-dependent PASEF, comprising 135,671 target precursors and 9,140 target proteins. For sample amounts on column of at least 200 ng and 120 min LC-MS runs, we reasoned that a diaPASEF method with a somewhat lower duty cycle, but higher precursor selectivity should be beneficial. We devised a method with four windows in each 100 ms diaPASEF scan. Eight of these scans covered the diagonal scan line for doubly charged peptides in the *m/z*-ion mobility plane and added a second, parallel scan line to ensure coverage of triply charged species with narrow 25 *m/z* precursor isolation windows.

For this acquisition scheme, the theoretical coverage of all library precursor ions was 97% and 95% for doubly and triply charged peptides, respectively (**Supplementary Fig. 2**).

In triplicate runs we detected a total of 80,545 peptide precursors (with 1% precursor and protein FDR), and on average 67,282 peptide precursors per run (**Fig. 4a**). The ion mobility values of precursors and fragment ions in the diaPASEF runs were highly correlated with the library values (r > 0.99, **Fig. 4b**), demonstrating the very high reproducibility of TIMS ion mobility values. The median absolute deviation of the fragment ion mobility values in diaPASEF to the library runs was 0.6% (**Fig. 4c**), the median summed absolute fragment mass deviation was 6.6 ppm and the median absolute retention time deviation was 17 s, which is 0.2% of the total LC-MS runtime. The combination of these three values defines the precision of the position of each precursor and its fragments in the diaPASEF data cuboid.

**Figure 4.**
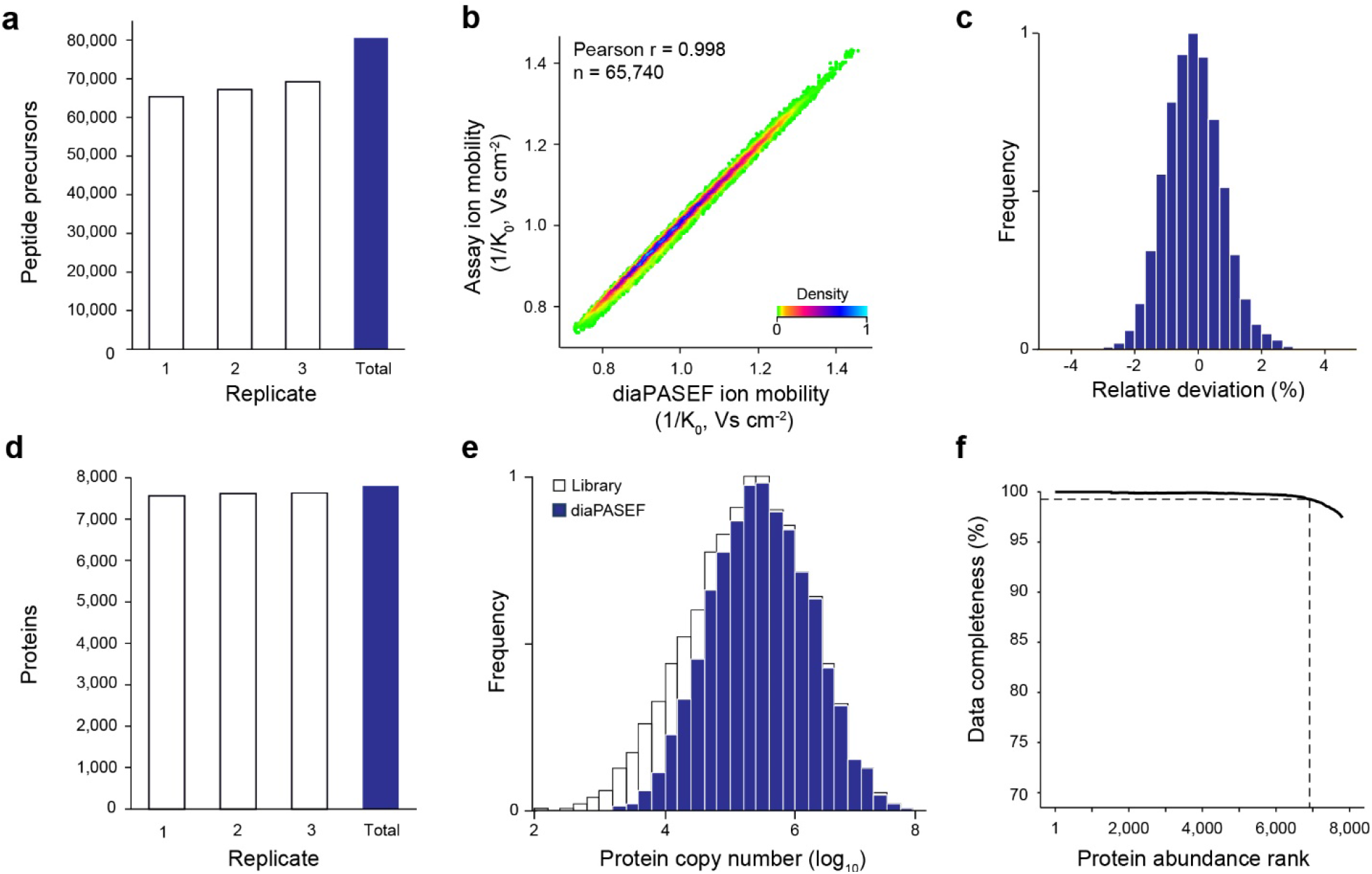
HeLa proteome analysis with diaPASEF. **a**Number of peptide precursor ions in triplicate injections of 200 ng HeLa digest. **b**, Correlation of precursor ion mobility in a single diaPASEF run and the assay library. **c**, Relative deviation of ion mobility values in a single diaPASEF run to the precursor ion mobility in the library. **d**, Number of proteins inferred from the peptide precursors in a. **e**, Distribution of estimated copy numbers of proteins contained in the assay library and detected with diaPASEF in triplicate single runs. **f,** Completeness of the protein quantification matrix in triplicate single runs as a function of decreasing protein abundance.

Overall, we identified 66,975 unique peptide sequences at 1% FDR, from which we inferred, on average, 7,600 proteins per run and 7,799 proteins in total at a global protein FDR of 1% (**Fig. 4d**). Proteins were inferred directly using only proteotypic peptides as mapped in the low-redundancy SwissProt database. In total, we covered a remarkable 85% of all proteins in the library in our 120 min single runs without fractionation. The quantified proteins spanned a dynamic range of about four orders of magnitude, as estimated by protein copy numbers derived from the library (**Fig. 4e**). Out of these, 7,347 proteins (94%) were quantified in all three replicates, 307 in two and only 145 proteins in a single replicate. Notably, the data completeness only dropped below 99% with the 10% least abundant proteins (**Fig. 4f**). The median coefficient of variation (CV) for fragment-ion based quantification was 10.4% on the protein level after median normalization.

### Label-free quantification benchmark

To benchmark the quantitative accuracy of diaPASEF in more detail, we set up a two-proteome experiment. We spiked 200 ng HeLa samples with approximately 45 ng and 15 ng of a tryptic yeast digest, respectively, and sequentially measured both samples in triplicate single runs as above. Mobi-DIK analysis using a combined human and yeast library quantified a total of 87,555 human and 7,603 yeast peptide precursors, from which we inferred 6,883 human and 1,300 yeast proteins. To compare the observed and expected abundance patterns of the identified proteins we used the LFQbench R package to evaluate our results consistently with previous reports^28^. Although the low-abundant yeast spike-in constituted only 7% of the sample, our stringent filtering left us with quantitative ratios for 6,778 human and 708 yeast proteins. The corresponding protein abundance ratios split in two distinct populations according to the mixing ratios (2.8 fold, **Fig. 5**). In line with the quantitative precision demonstrated above, the human population clustered precisely around the 1:1 ratio throughout the full abundance range (σ(log_2_) = 0.29). The low-abundance yeast spike-ins were quantified with a somewhat lower overall precision (σ(log_2_) = 0.67), yet quantitatively similar to human proteins in the same abundance range. We thus conclude that our label-free diaPASEF workflow precisely and accurately quantifies changes in protein abundance.

**Figure 5.**
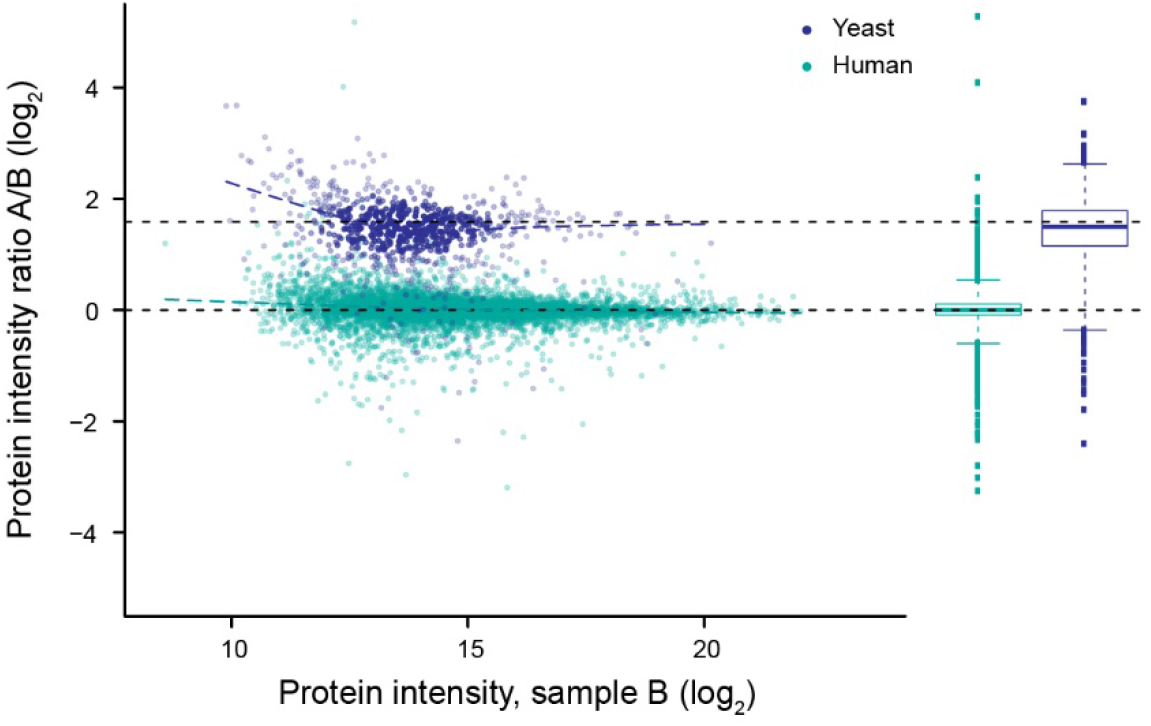
Label-free protein quantification benchmark. HeLa digest was spiked with approximately 45 ng (sample A) and 15 ng (sample B) yeast digest and analyzed in triplicate diaPASEF single runs each. Log-transformed ratios are plotted as a function of protein abundance for n = 6,777 human and n = 708 yeast proteins. Boxplots show the median (center line), 25^th^ and 75^th^ percentile (lower and upper box limits), the 1.5 × the interquartile range (whiskers) and outliers (points).

### Very high sensitivity proteomics with diaPASEF

One of the strengths of diaPASEF is that it can be tuned to utilize a very high fraction of the incoming ion beam and still achieve a high precursor selectivity. This is because single-charge background ions are excluded from the analysis and because TIMS separates chromatographically co-eluting precursors present in the same mass window. One application of this principle is the accumulation of high ion signal for low-abundance precursors. To demonstrate this concept, we analyzed only 10 ng of HeLa digest in triplicate 120 min single runs. We employed a diaPASEF scheme that samples about 25% of the ion current and covers the most dense precursor ion region (**Supplementary Fig. 3**). This high duty cycle increased the detected fragment ion signal on average about 4-fold as compared with the standard diaPASEF method used above (**Fig. 6a**). The higher ion signal translated into a more precise quantification for the same set of peptides, in particular for low-abundance peptide precursors. Even though the high-duty cycle method covers a narrower precursor space in *m/z*-ion mobility dimensions, we quantified on average about 13,000 peptides with each method. In effect, the high-duty cycle extends the lower limit of detection by about 4-fold, which is almost directly proportional to the raw signal increase (**Fig. 6b**). The standard diaPASEF method already quantified on average 3,537 proteins per injection of 10 ng HeLa digest, highlighting the intrinsic high sensitivity of diaPASEF and the TIMS-QTOF setup. The high-duty cycle method further increased this to 3,785 proteins on average. Cumulatively, we quantified 4,250 proteins in triplicates of 10 ng injections with a data completeness of 89% (**Fig. 6c**). However, at higher sample amounts of 100 ng narrower quadrupole windows are more beneficial.

**Figure 6.**
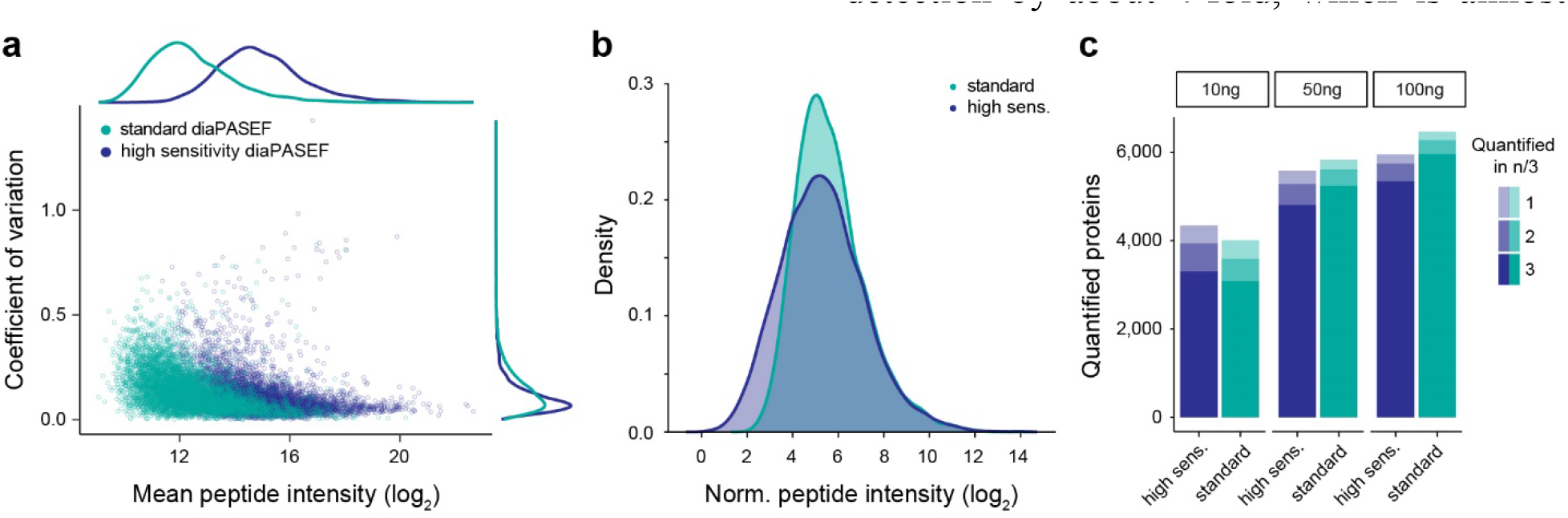
High duty cycle diaPASEF analysis of low sample amounts. **a,** Summed fragment ion intensity and coefficient of variation of shared peptides in triplicate injections of 10 ng HeLa digest with different diaPASEF duty cycles. **b**, Total peptide intensity distribution for both methods. **c**, Quantified proteins in n of 3 replicate injections of 10, 50 and 100 ng HeLa digest.

## DISCUSSION

Here we have developed and demonstrated a PASEF workflow in a TIMS-TOF mass spectrometer that implements the DIA principle. Making use of the correlation between the ion mobility and the *m/z* of peptides, precursors are trapped and then released in sync with the quadrupole position in our diaPASEF scheme, resulting in almost complete sampling of the precursor ion beam. This is in contrast to DDA methods, which convert only a very small fraction – generally much less than 1% - of the incoming ion beam into fragments and even typical DIA workflows, which convert a few percent of the ion beam at best. On low complex mixtures, we achieve close to 100% of the theoretical maximum, whereas for more complex mixtures, it was beneficial to use the quadrupole to decrease spectral complexity and increase selectivity, thereby somewhat reducing the fraction of total available ions sampled. To extract information using spectral library-based targeted data analysis, we extended the OpenSWATH tool developed for DIA applications to efficiently make use of the ion mobility dimension for library matching, providing full FDR control and excellent quantification.

Even in this first implementation, we achieved deep proteome coverage of more than 7,000 proteins in single, 2 h LC runs from a total of 200 ng HeLa peptide sample on column with high degree of reproducibility. Our two-proteome experiment verifies that the quantitative accuracy of the method is in line with previous strategies even when substantially constrained by the lower loading amount of yeast (15 ng). Even more remarkably, we detected over 4,000 proteins in triplicate injections of only 10 ng HeLa peptide mass on column. This latter result points to a perhaps unexpected advantage of diaPASEF, namely the fact that the high ion sampling also fully translates to higher sensitivity. Likewise, the very fast cycle time of our new scan mode should be very advantageous for short gradients, an increasingly important attribute as large scale biological and clinical studies require very large throughput. For the future, we imagine that both hardware and software can still be greatly optimized to further increase the amount and quality of the information contained in and extracted from the extremely rich four-dimensional diaPASEF data cuboids. Furthermore, we note that applications of diaPASEF are not restricted to peptides but could equally well be extended to metabolites, lipids or other compound classes^21^.

## METHODS

### Sample preparation

The human cancer cell line (HeLa S3, ATCC) was cultured in Dulbecco’s modified Eagle’s medium with 10% fetal bovine serum, 20 mM glutamine and 1% penicillin-streptomycin. Cells were collected by centrifugation, washed with phosphate-buffered saline, flash-frozen in liquid nitrogen and stored at −80°C. Cell lysis, reduction and alkylation was performed in lysis buffer with chloroacetamide (PreOmics) following our previously published protocol^29^. Briefly, the cell suspension was heated to 95°C for 10 min and subsequently sonificated to further disrupt cells and shear nucleic acids. Proteins were enzymatically cleaved overnight by adding equal amounts of LysC and trypsin in a 1:100 (wt/wt) enzyme:protein ratio. De-salting and purification was performed according to the PreOmics iST protocol on a styrendivinylbenzene reversed-phase sulfonate (SDB-RPS) sorbent. Purified peptides were vacuum-centrifuged to dryness and reconstituted in double-distilled water with 2 vol.-% acetonitrile (ACN) and 0.1 vol.-% trifluoroacetic acids (TFA) for single run LC-MS analysis or fractionation.

To evaluate the quantitative accuracy of diaPASEF, we performed a two-proteome experiment with HeLa and yeast. Purified and predigested yeast standard was purchased from Promega (Madison, USA) and resuspended in 0.1% formic acid (FA). Whole HeLa cell pellets were purchased from CIL Biotech (Mons, Belgium). Cell lysis was performed using trifluoroethanol (TFE)^30^. Briefly, the cell suspension was kept 10 minutes on ice and then incubated at 56°C for 20 min. We used 200 mM dithiothreitol (DTT) to reduce proteins at 90°C for 20 min and 200mM iodoacetamide (IAA) to alkylate free cysteine residues (90 min at room temperature). Proteins were enzymatically cleaved overnight by adding trypsin in a 1:100 (wt/wt) enzyme:protein ratio. De-salting and purification was done using a solid phase extraction cartridge (Empore C_18_ SPE cartridge, Sigma Aldrich, St. Louis, USA) by diluting and washing protein digest with 0.1% FA and subsequent elution with 50% (w/w) ACN in 0.1% FA. Purified and dried peptides were reconstituted in 0.1% FA. For the two-proteome experiment, the purified peptides from HeLa and yeast were combined as following: Sample A was composed of 200 ng human and 45 ng yeast proteins per LC-MS injection, and sample B of 200 ng human and 15 ng yeast proteins per LC-MS injection.

### High-pH reversed-phase fractionation

To generate a comprehensive library of HeLa precursor and fragment ions, peptides were fractionated at pH 10 with a ‘spider fractionator’ coupled to an EASY-nLC 1000 chromatography system (Thermo Fisher Scientific) as described previously^31^. Approximately 50 μg purified peptides were separated on a 30 cm C_18_ column within 96 min and automatically concatenated into 24 fractions by shifting the exit valve every 120 s. The fractions were vacuum-centrifuged to dryness and re-constituted in double-distilled water with 2 vol.-% ACN and 0.1 vol.-% trifluoroacetic acids TFA for LC-MS analysis. To generate a library for the two-proteome experiment, 100 μg purified peptides from both yeast and HeLa digest were each fractionated at pH 10 on a reversed phase column (Waters Acquity CSH C18 1.7 μm 1 × 150 mm) using a Dionex Ultimate 3000 system (Thermo Fisher Scientific). The fractions were freeze-dried and re-constituted in 0.1% formic acid.

### Liquid chromatography

Nanoflow reversed-phase chromatography was performed on an EASY-nLC 1200 system (Thermo Fisher Scientific). Peptides were separated within 120 min at a flow rate of 300 nL/min on a 50 cm × 75 μm column with a laser-pulled electrospray emitter packed with 1.9 μm ReproSil-Pur C_18_-AQ particles (Dr. Maisch). Mobile phases A and B were water with 0.1 vol.-% formic acid and 80/20/0.1 vol.-% ACN/water/formic acid. The %B was linearly increased from 5 to 30% within 95 min, followed by an increase to 60% within 5 min and a further increase to 95% before re-equilibration. For the two-proteome experiment, we employed a nanoElute liquid chromatography system (Bruker Daltonics). Peptides were separated within 120 min at a flow rate of 400 nL/min on a commercially available reversed-phase C_18_ column with an integrated CaptiveSpray Emitter (25 cm × 75μm, 1.6 μm, IonOpticks, Australia). Mobile phases A and B were with 0.1 vol.-% formic acid in water and 0.1% formic acid in ACN. The fraction of B was linearly increased from 2 to 25% within 90 min, followed by an increase to 35% within 10 min and a further increase to 80% before re-equilibration.

### Mass spectrometry

LC was coupled online to a hybrid TIMS quadrupole time-of-flight mass spectrometer (Bruker timsTOF Pro) via a CaptiveSpray nano-electrospray ion source. A detailed description of the instrument is available in ref. ^20^. The dual TIMS analyzer was operated at a fixed duty cycle close to 100% using equal accumulation and ramp times of 100 ms each. We performed data-dependent data acquisition in PASEF mode with 10 PASEF scans per topN acquisition cycle. Singly charged precursors were excluded by their position in the *m/z*-ion mobility plane and precursors that reached a ‘target value’ of 20,000 a.u. were dynamically excluded for 0.4 min. The quadrupole isolation width was set to 2 Th for *m/z* < 700 and 3 Th for *m/z* > 700.

To perform data-independent acquisition, we extended the instrument control software to define quadrupole isolation windows as a function of the TIMS scan time (diaPASEF). The instrument control electronics were modified to allow seamless and synchronous ramping of all applied voltages. We tested multiple schemes for data-independent precursor windows and placement in the *m/z*-ion mobility plane and defined up to 8 windows for single 100 ms TIMS scans as detailed in the main text. Acquisition schemes for the three diaPASEF methods used herein are shown in **Supplementary Figs. 1-3**. In both scan modes, the collision energy was ramped linearly as a function of the mobility from 59 eV at 1/K_0_=1.6 Vs cm^−2^ to 20 eV at 1/K_0_=0.6 Vs cm^−2^.

### Spectral library generation

To generate spectral libraries for targeted data extraction, we first analyzed the high pH reversed-phase fraction acquired in DDA mode with MaxQuant version 1.6.5.0, which extracts four-dimensional features on the MS1 level (retention time, m/z, ion mobility and intensity) and links them to peptide spectrum matches. The maximum precursor mass tolerance of the main search was set to 20 ppm and deisotoping of fragment ions was deactivated. Other than that, we used the default ‘TIMS-DDA’ parameters. MS/MS spectra were matched against an *in silico* digest of the Swiss-Prot reference proteome (human 20,414 entries, yeast 6,721 entries, downloaded July 2019) and a list of common contaminants. The minimum peptide length was set to 7 amino acids and the peptide mass was limited to 4,600 Da. Carbamidomethylation of cysteine residues was defined as a fixed modification, methionine oxidation and acetylation of protein N-termini were defined as variable modifications. The false discovery rate was controlled <1% at both, the peptide spectrum match and the protein level. Our Mobi-DIK software package builds on OpenMS tools to compile spectral libraries in the standardized TraML or pqp formats from the MaxQuant output tables while retaining the full ion mobility information for each precursor-to-fragment ion transition. Only proteotypic peptides with a precursor *m/z* >400 were included in the library and required to have a minimum of 6 fragment ions with *m/z* >350 and outside the precursor mass isolation range.

### Targeted data extraction

To analyze diaPASEF data, we developed an ion mobility DIA analysis kit (Mobi-DIK) that extracts fragment ion traces from the four-dimensional data space as detailed in the main text. Raw data were automatically re-calibrated using curated reference values in *m/z*, retention time and ion mobility dimensions (387 peptides for linear and 3,184 peptides for non-linear alignment). We applied an outlier detection in each dimension before calculating the final fit function to increase robustness. Peak picking and sub-sequent scoring functionalities in the Mobi-DIK software build on OpenSWATH modules. For diaPASEF, we extended these modules to also take into account the additional ion mobility dimension. OpenSWATH (Revision: e0b987a) was run with following parameters: min_coverage = 0.1, RTNormalization:alignmentMethod = lowess, RTNormalization:lowess:span = 0.01, Scoring:TransitionGroupPicker:PeakPicker MRM:sgolay_frame_length = 11, Scoring:stop_report_after_feature = 5, rt_extraction_window = 250, Scoring:Scores:use_ion_mobility_scores, mz_correction_function = quadratic_regression_delta_ppm, use_ms1_traces, mz_extraction_window = 25, mz_extraction_window_unit = ppm, mz_extraction_window_ms1 = 25, mz_extraction_window_ms1_unit = ppm, irt_mz_extraction_window_unit = ppm, irt_mz_extraction_window = 40, Calibration:ms1_im_calibration, ion_mobility_window = 0.06, irt_im_extraction_window = 99, RTNormalization:NrRTBins = 8, RTNormalization:MinBinsFilled = 4. All other parameters were set to default. PyProphet was used to train an XGBoost classifier for target-decoy separation by first creating one concatenated and sub-sampled OpenSwath output for each set of three replicate injections of the same acquisition strategy and sample amount. The classifier was subsequently applied for scoring all samples, controlling the FDRs <1% at the peak group level per sample and at both global peptide and global protein level. In case of two overlapping diaPASEF windows, the analysis was performed separately for the individual windows and for FDR estimations, the highest scoring peak group was selected. Protein abundances were inferred using the ‘best flier peptide’ approach summing the intensities of the top 5 fragment ions from the top 3 peptides^32,33^. Potential contaminants were excluded from further analysis.

### Bioinformatics

Output tables from the data analysis pipeline or MS raw data were further analyzed and visualized in the R statistical computing environment or in Python v3.6. Protein copy numbers were estimated with the Proteomic Ruler^34^ Perseus^35^ plugin. We used the LFQbench^28^ R package to analyze the two-proteome experiment.

## Data availability

The mass spectrometry datasets generated during and analyzed during the current study have been deposited to the ProteomeXchange Consortium via the PRIDE^36^ partner repository with the dataset identifier: PXD017703 (user, password: LWudexAH).

## Code availability

Code is available under the three-clause BSD license on https://github.com/OpenMS/OpenMS and https://github.com/Roestlab/dia-pasef.

## ACKNOWLEDGEMENTS

This work was partially supported by the German Research Foundation (DFG-Gottfried Wilhelm Leibniz Prize granted to M.M.) and by the Max Planck Society for the Advancement of Science. This work was partially supported by the Government of Canada through Genome Canada (grant no. 15411) and the Canadian Institutes for Health Research. B.C. was supported by a Swiss National Science Foundation Ambizione grant (no. PZ00P3_161435). R.A. was supported by the Swiss National Science Foundation (grant no. 3100A0-688 107679) and the European Research Council (ERC-20140AdG 670821). We thank our colleagues in the department of Proteomics and Signal Transduction and at Bruker Daltonik for discussion and help, in particular J. Müller, A. Strasser, C. Deiml and I. Paron for technical support.

## AUTHOR CONTRIBUTIONS

F.M., R.A., B.C., H.R. and M.M. conceptualized and designed the study; F.M. and M.M. conceived the acquisition mode; H.R. conceived the data analysis software; F.M., A.B., S.K., M.L., O.R. and B.C. performed experiments; A.H. and M.F. contributed to the software development; F.M., A.B., M.F., A.H., I.B., E.V., S.K., B.C., H.R. and M.M. analyzed the data; F.M., R.A., B.C., H.R. and M.M. wrote the manuscript.

