## Supplementary Information for "Parallel accumulation – serial fragmentation combined with data-independent acquisition (diaPASEF): Bottom-up proteomics with near optimal ion usage"

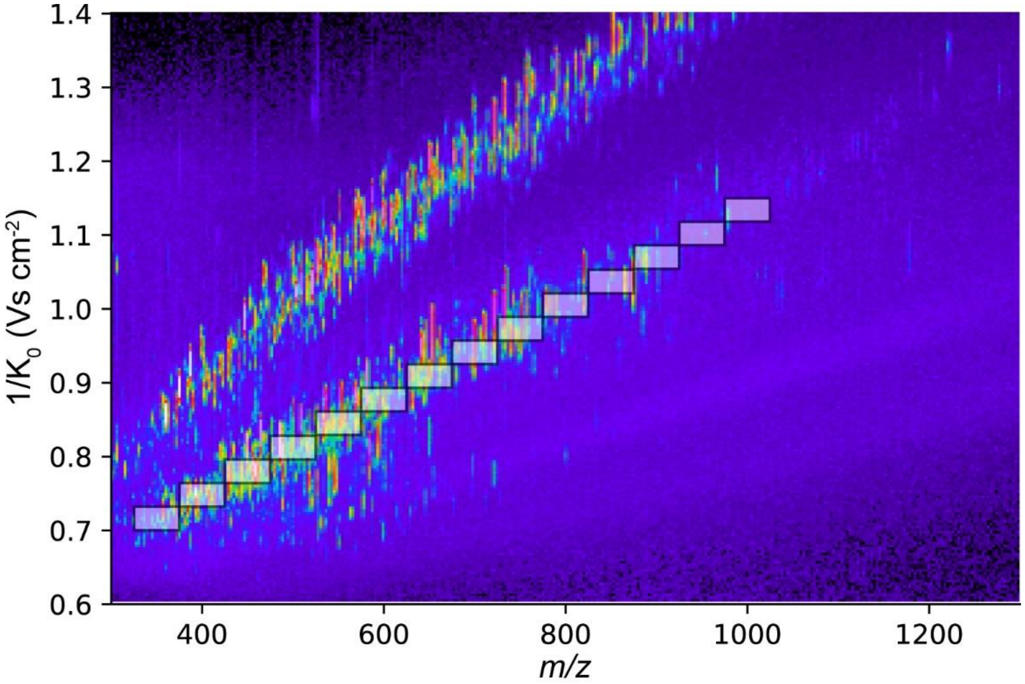


**Supplementary Figure 1.** Position of the precursor isolation windows in a diaPASEF acquisition scheme with close to 100% duty cycle overlaid on the average precursor ion intensity in a 45 min LC-MS experiment of BSA digest.

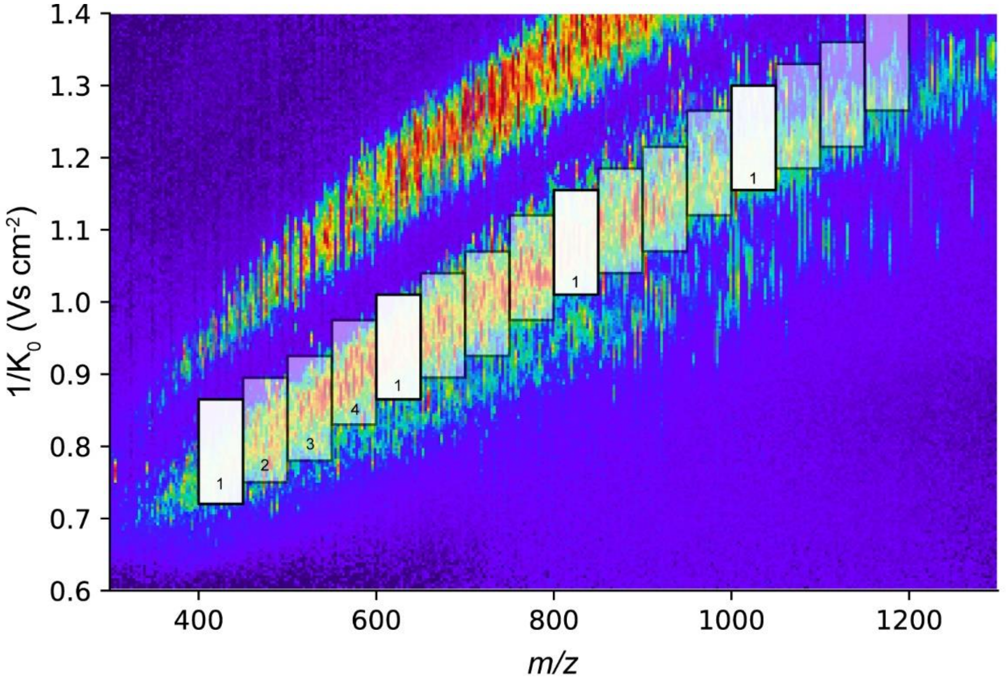


**Supplementary Figure 2.** Position of the precursor isolation windows in a diaPASEF acquisition scheme with 4 scans overlaid on the average precursor ion intensity in a 120 min LC-MS experiment of HeLa digest. Numbers indicate individual diaPASEF scans (first diaPASEF scan highlighted).

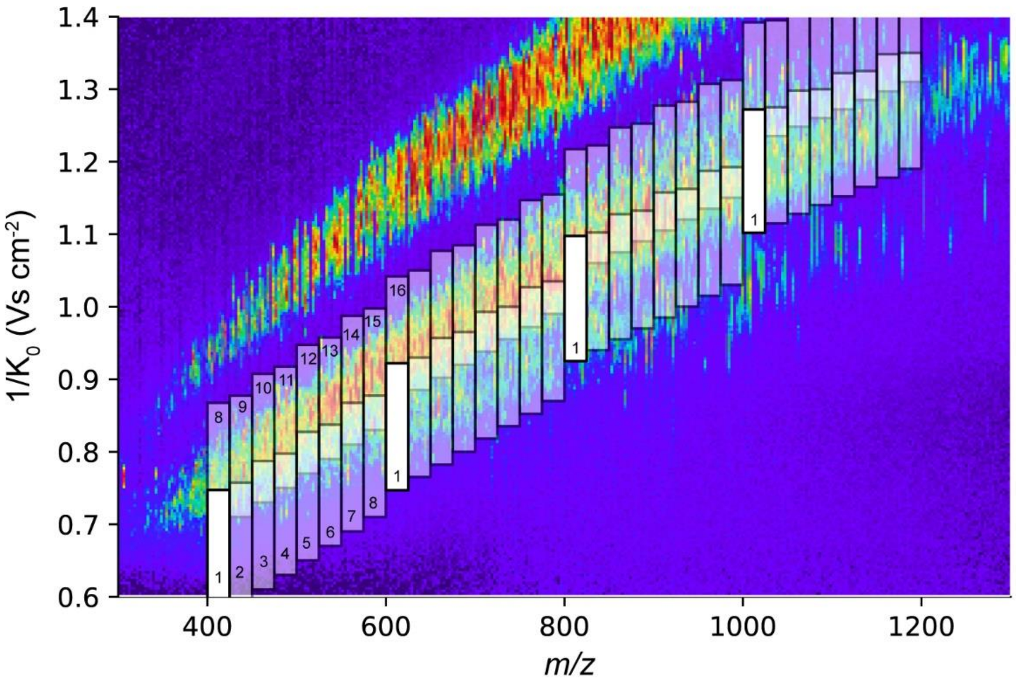


**Supplementary Figure 3.** Position of the precursor isolation windows in a diaPASEF acquisition scheme with 16 scans overlaid on the average precursor ion intensity in a 120 min LC-MS experiment of HeLa digest. Numbers indicate individual diaPASEF scans (first diaPASEF scan highlighted).
